# Interpretable deep learning of myelin histopathology in age-related cognitive impairment

**DOI:** 10.1101/2022.06.06.495016

**Authors:** Andrew T. McKenzie, Gabriel Marx, Daniel Koenigsberg, Mary Sawyer, Megan A. Iida, Jamie M. Walker, Timothy E. Richardson, Gabriele Campanella, Johannes Attems, Ann C. McKee, Thor D. Stein, Thomas J. Fuchs, Charles L. White, The PART working group, Kurt Farrell, John F. Crary

## Abstract

Age-related cognitive impairment is multifactorial, with numerous underlying and frequently co-morbid pathological correlates. Amyloid beta (Aβ) plays a major role in Alzheimer’s type age-related cognitive impairment, in addition to other etiopathologies such as Aβ-independent hyperphosphorylated tau, cerebrovascular disease, and myelin damage, which also warrant further investigation. Classical methods, even in the setting of the gold standard of postmortem brain assessment, involve semi-quantitative ordinal staging systems that often correlate poorly with clinical outcomes, due to imperfect cognitive measurements and preconceived notions regarding the neuropathologic features that should be chosen for study. Improved approaches are needed to identify histopathological changes correlated with cognition in an unbiased way. We used a weakly supervised multiple instance learning algorithm on whole slide images of human brain autopsy tissue sections from a group of elderly donors to predict the presence or absence of cognitive impairment (*n* = 367 with cognitive impairment, *n =* 349 without). Attention analysis allowed us to pinpoint the underlying subregional architecture and cellular features that the models used for the prediction in both brain regions studied, the medial temporal lobe and frontal cortex. Despite noisy labels of cognition, our trained models were able to predict the presence of cognitive impairment with a modest accuracy that was significantly greater than chance. Attention-based interpretation studies of the features most associated with cognitive impairment in the top performing models suggest that they identified myelin pallor in the white matter. Our results demonstrate a scalable platform with interpretable deep learning to identify unexpected aspects of pathology in cognitive impairment that can be translated to the study of other neurobiological disorders.

## Introduction

Cognitive impairment is not an invariable part of the aging process and unimpaired cognition is a core feature of most criteria of successful aging [1]. While Alzheimer’s disease (AD) type amyloid-beta peptide (Aβ) deposition in senile plaques may play a role in age-related cognitive impairment, it is clear that removing or ameliorating Aβ alone will not alleviate all cognitive impairment in aging [2]. The neuropathologic correlates of cognitive impairment are multifactorial, with mixed pathologies accounting for the majority of cases in community samples [3, 4]. Data suggest that multiple forms of brain pathology can each be uniquely associated with risk of age-related cognitive impairment, including cerebrovascular disease, neuritic plaques and neurofibrillary tangles, Lewy body disease, TDP-43 pathology, and hippocampal sclerosis [5, 6]. To pave the way towards better prevention and treatment options for age-related cognitive impairment, there is an urgent need to identify the structural features of brain microanatomy that are robustly associated with the condition using unbiased assessment protocols [7, 8]. One approach to identifying structural correlates of cognitive impairment is to perform clinicopathologic correlation in postmortem human brains.

Recent advances in digital pathology, namely whole slide image (WSI) scanning and analysis, provide an opportunity to address the question of clinicopathologic correlation in a way that is less biased towards established paradigms [9]. Studies have begun to apply computational analysis of WSI data using deep learning to answer neuropathologic questions [10–12]. However, the use of deep learning in neuropathology has often been limited by the need for intensive manual annotations. Moreover, deep learning analysis in neuropathology has often used supervised learning to study an existing domain of structural features, rather than the discovery of potentially unexpected features.

Weakly supervised deep learning offers a clear path towards WSI analysis in neuropathology with less bias and without the need for laborious manual annotations. In weakly supervised learning, the deep learning algorithm attempts to classify the WSI on the basis of a single slide-level diagnosis or label, rather than pixel-level inputs [13]. Weakly supervised learning approaches using multiple instance learning have had remarkable success thus far in digital pathology, especially in oncology [13, 14].

However, unlike cancer pathology, where a gold standard diagnosis can be ascertained, the neuropathologic etiologies of cognitive impairment are poorly understood, graded rather than categorical, overlapping, and dynamically interacting [15–17]. Moreover, clinical measures of cognitive function available for clinicopathologic correlation in neuropathology are frequently imprecise, non-standardized, ephemeral, and collected at distant time points prior to death [18–20]. As a result, the use of weakly supervised learning to correlate age-related cognitive impairment with neuropathologic features using WSI data will be dependent on noisy labels of cognition.

To the best of our knowledge, no study has yet reported a weakly supervised deep learning approach on brain tissue WSI data to identify features associated with age-related cognitive impairment. It is uncertain the degree to which deep learning models will be able to identify robust features to make the prediction of whether an autopsy brain donor had antemortem cognitive impairment in the setting of noisy labels. In this study, we used WSI data stained with Luxol fast blue (LFB), hematoxylin, and eosin (LH&E), from the hippocampus and frontal cortex in a previously described cohort of elderly individuals with a spectrum of age-related pathologies [21–25]. We leveraged a published weakly supervised deep learning algorithm, clustering-constrained-attention multiple instance learning (CLAM) [14], on this histopathologic data to identify pathoanatomic features that are associated with cognition. Our approach re-purposes the classification procedure as a method for inferring pathoanatomical group differences between those found to have any aspect of cognitive impairment and those who were not. We explored the association between the deep learning model predictions on neuropathologic data and antemortem evidence of cognitive impairment. To interpret these results, we dissected the deep learning model’s attention weights using additional machine vision techniques. Our study shows that weakly supervised deep histopathology is a promising platform to perform clinicopathologic correlation in neuropathology.

## Materials and methods

### Description of the overall cohort and subset analyzed in this study

Our study used digital WSIs of stained formalin-fixed paraffin embedded (FFPE) tissue from the frontal cortex and hippocampus of a subset of individuals from a previously described collection [21–23]. The cohort is a convenience sample derived from our ongoing studies of brain aging, which was collected by eliciting samples from multiple institutions. Extensive neuropathological assessments were completed at the contributing institutions using standardized criteria. This assessment included CERAD neuritic plaque severity score and Braak stage [26]. This cohort contains individuals with varying degrees of primary age-related tauopathy (PART) pathologic change, including PART possible (mild amyloid plaques) and PART definite (amyloid plaque negative), among other age-related changes [27]. Neuropathological exclusion criteria consisted of other neurodegenerative diseases including Lewy body disease, progressive supranuclear palsy (PSP), corticobasal degeneration (CBD), chronic traumatic encephalopathy (CTE), Pick disease, Guam amyotrophic lateral-sclerosis-parkinsonism-dementia, subacute sclerosing panencephalitis, globular glial tauopathy, and hippocampal sclerosis. There are also individuals that do not meet the neuropathologic criteria for PART (*e.g.*, two cases with moderate amyloid), and therefore it should be considered an aging-related cognitive impairment cohort. Cerebrovascular pathology was defined in an inclusive manner based on clinical or gross pathoanatomic evidence of vascular disease in the brain in the provided records. In this cohort, ARTAG positivity or absence was assessed on matched phosphorylated tau immunohistochemical stains as previously described [22].

Inclusion criteria in the subset of this cohort analyzed in this paper were individuals who had antemortem clinical evidence of either normal cognition or cognitive impairment, while those without such data were excluded. This led to a data set with WSI and matched pathoclinical data from a total of *n* = 716 donors (**Table 1**). For the definition of cognitive impairment, we used a hierarchical method based on the three metrics in the available clinical data to identify any evidence of cognitive impairment. First, if available, a clinical dementia rating (CDR) score >= 0.5 was used as the primary measure of cognitive impairment; if CDR was not available, then the presence of any clinical diagnosis suggestive of cognitive impairment was used as the secondary measure; and finally, if the first two more global metrics were not available, then a Mini-Mental State Examination (MMSE) score <= 24 was used as a measure of cognitive impairment [28]. To maximize the sample size available for the study, cognitive data was included even if the time of assessment relative to death was unknown. Brain donors with any evidence of cognitive impairment were considered a part of the cognitively impaired (CI) group, while donors with negative data in all the available categories were included in the non-cognitively impaired (NCI) group. CDR scores with dementia severity score greater than 3 were converted to a maximum of 3 for consistency across centers.

**Table 1.** Description of cohort subset and whole slide image dataset used in this study.

|  |  |  |  |  |
| --- | --- | --- | --- | --- |
| **Category** | **Non-cognitively impaired** | **Cognitively impaired** | **Total group** | **p-value for difference\*** |
| **Sample size** | 367 | 349 | 716 | Not applicable |
| **Age (Mean +/- SEM)** | 83.0 +/- 0.58 | 87.4 +/- 0.48 | 85.2 +/- 0.39 | $p = 1.02 \times 10^{-8}$ |
| **Proportion female** | 0.54 | 0.53 | 0.54 | *Not significant* |
| **Mean Braak score** | 2.4 | 2.5 | 2.4 | *Not significant* |
| **Proportion CERAD neuropathology positive** | 0.15 | 0.18 | 0.17 | *Not significant* |
| **Proportion hippocampal ARTAG positive** | 0.22 | 0.31 | 0.27 | $p = 0.015$ |
| **Proportion with hippocampal WSI** | 0.99 | 0.99 | 0.99 | *Not significant* |
| **Proportion with frontal WSI** | 0.46 | 0.46 | 0.46 | *Not significant* |

This table describes the pathoclinical characteristics of the subset of brain donors employed in this study. The significance of differences in categorical variables between the non-cognitively impaired and cognitively impaired groups was assessed with a two-proportions z-test, while the significance of differences in numerical variables was assessed with a t-test. WSI = Whole slide image; SEM = Standard error of the mean; ARTAG = Aging-Related Tau Astrogliopathy; CERAD = Consortium to Establish a Registry for Alzheimer's Disease.

### Slide preparation

Luxol fast blue, hematoxylin, and eosin (LH&E) stains were performed on 4-µm-thick FFPE sections as previously described [23]. Sections mounted on positively charged slides were dried overnight. For each batch of slides stained, a known severe AD case was included as a positive staining control. WSI were scanned using an Aperio CS2 (Leica Biosystems, Buffalo Grove, IL) digital slide scanner at 20x magnification (0.5 microns per pixel).

### Weakly supervised learning pipeline

We used Python (v. 3.7.7), PyTorch (v. 1.3.1), and CLAM to perform deep learning on WSIs [14]. Models were trained using 4 NVIDIA V100 GPUs available on Minerva, a high-performance computing cluster at the Icahn School of Medicine at Mount Sinai. LH&E WSIs were segmented into tiles of 256 x 256 pixels using the default automated segmentation settings in CLAM. All the tissue in the WSI was included in the segmentation and downstream analysis. For example, for WSIs generated from blocks with two tissue sections on the slide, both tissue sections were automatically segmented and used in downstream analyses.

To perform feature extraction, for each tile, the first three blocks of a ResNet50 model pre-trained on ImageNet was used to convert each 256 x 256-pixel tile into a 1024-dimensional feature vector. Training in CLAM uses attention-based pooling to leverage tile-level feature vectors in assembling slide-level representations for each of the two classes. For network training, we used the default CLAM parameters, with a single attention branch model and a learning rate of 2e-4. For WSI classification training, we used 10-fold Monte Carlo cross-validation to split the data set into 10 folds of training sets (80% of cases), validation sets (10% of cases), and test sets (10% of cases). A separate model was trained on each of the 10 folds, using performance on the validation set for early stopping during training, and performance on the test set at the end of training as a measure of prediction accuracy.

We used R (v. 4.0.1) and ggplot2 (v. 3.3.5) to perform downstream analysis and visualization of results from the weakly supervised deep learning analysis. To evaluate the performance of the deep learning algorithm, we compared the performance of all 10 independently trained models to chance (*i.e.*, an area under the receiver operating characteristic (ROC) curve, or AUC, of 0.5) using one-sample Wilcoxon signed rank tests with continuity correction and plotted the average ROC curves using vertical averaging and linear interpolation [29]. For the analysis from each of the two brain regions studied, we used the best-performing model, as measured by the arithmetic mean of the area under the curve and the balanced accuracy on the test set, for subsequent analyses. To perform differential rank correlation analysis between groups, we used the DGCA package [30], with 10,000 permutations of the data used to generate empirical p-values.

### Attention interpretation analysis

For each WSI, we used CLAM to perform tissue-level attention analysis of the top performing trained models. In these heatmaps, the red colors represent regions assigned relatively high attention by the model and blue colors represent regions assigned relatively low attention, normalized to the attention values in the rest of the slide.

To evaluate the macrostructural features most associated with cognitive impairment, we used V7 to annotate the macroscopic tissue types in a randomly chosen subset of WSIs from both the hippocampus and frontal cortex. One trained researcher (M.S.) created the annotations, and an expert neuropathologist (J.F.C.) reviewed them to ensure accuracy. The V7 annotations were converted to the same tile-level space as the tile-level attention score output from CLAM. We z-transformed the attention scores and we then calculated the median tile-level attention score for each tissue region within each slide. We compared the median attention scores across tissue types with paired t-tests.

To evaluate the microstructural features most associated with cognitive impairment, we examined the 100 tiles with the highest attention scores from each WSI. In order to quantify the amount of dark blue staining in LHE stained tiles, we used the positive pixel counting function in the Python package HistomicsTK (v 0.1.0) [31]. This function converts RGB color space images to HSI (hue, saturation, intensity) color space and calculates the number of pixels in a user-defined hue range. The parameters used to count the positive pixels were created based on manually identifying the appropriate dark blue hue range in HSI space (**Supplementary Fig 1**). As a normalization measure, we also measured the ratio of the dark blue to light blue color stain in each tile. Outlier tiles with zero positive pixels were removed from further analysis. The same positive pixel counting analysis pipeline was applied to each of the top 100 attention tiles identified from each WSI in the data sets. To minimize the impact of outliers, the median of the results was found for each slide. The slide-level median values between different groups were then compared with t-tests. The multivariate combination of the total dark blue pixel counts and the ratio of dark to light blue pixel counts between groups predicted to be cognitively impaired or not were compared with two-dimensional kernel density estimation using the MASS R package (v. 7.3-51.6) and visualized with contour lines.

**Figure 1.**
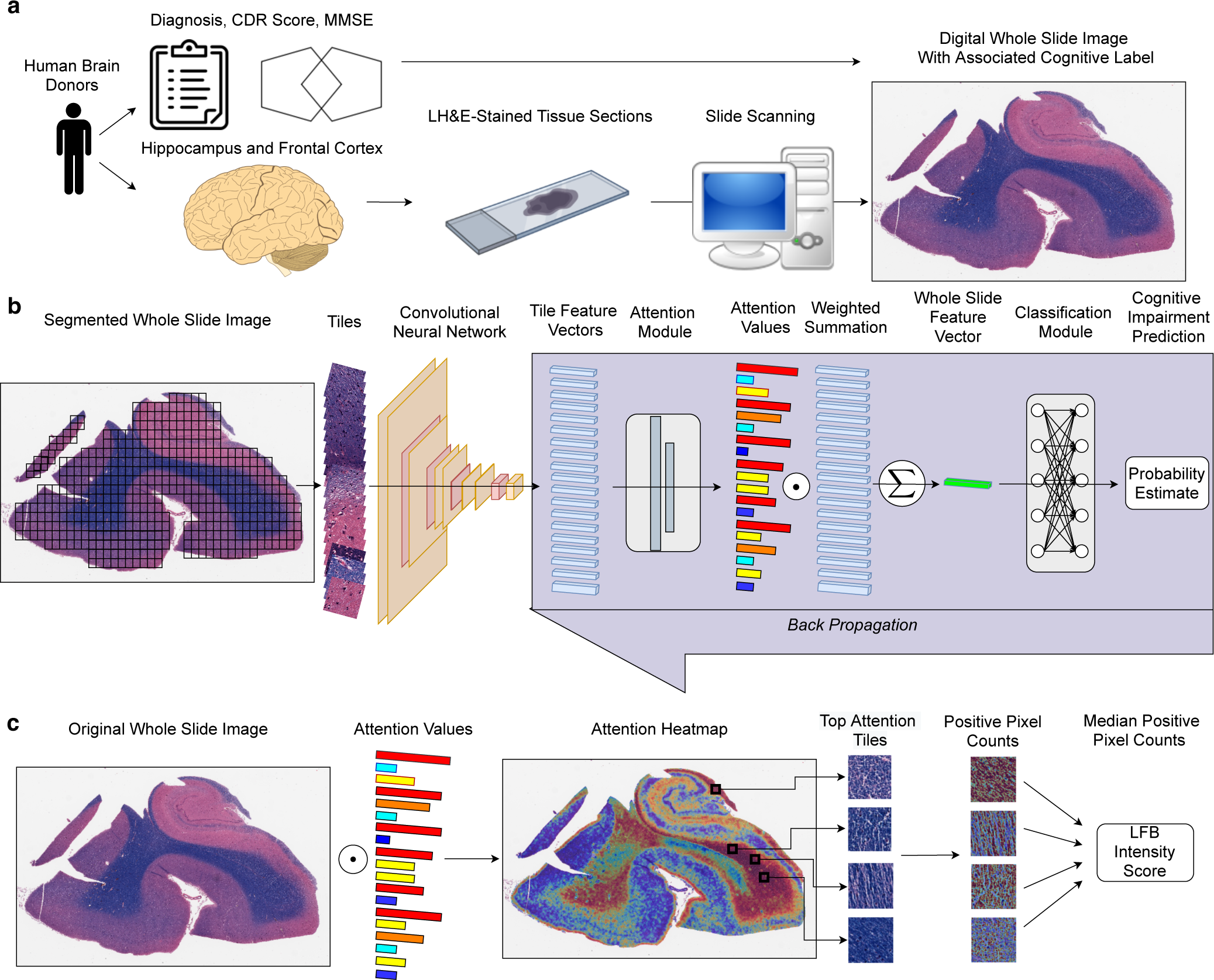
Workflow for performing weakly supervised deep learning of age-related cognitive impairment. **A**: Generation of digital neuropathology whole slide images (WSI) with associated cognitive labels. Human brain sections were stained with Luxol fast blue (LFB) and counterstained with hematoxylin & eosin (LH&E). Cognitive labels were generated based on clinical diagnosis, clinical dementia rating (CDR) scores, and/or mini-mental state exam (MMSE) scores. **B**: WSI were segmented into tiles and passed through a convolutional neural network for feature extraction. The resulting tile-level feature vectors were passed through an attention network. Each feature vector was multiplied by its associated attention score and a weighted summation operation was performed to create slide-level feature vectors. The slide-level feature vectors were then passed through a classification network. The attention and classification networks were trained via backpropagation. **C**: For interpretation analysis, attention heatmaps were created by mapping the attention scores at their associated tile locations in the original WSI. Among the top attention tiles, a dark blue hue range associated with LFB staining was counted and quantified to calculate a slide-level median staining intensity value.

### Association of deep learning predictions with pathoclinical traits

We used rank correlation analysis to compare slide-level probability estimates of cognitive impairment and slide-level averages of dark blue color density with other pathoclinical traits in the data set, namely age, cerebrovascular pathology, hippocampal aging-related tau astrogliopathy (ARTAG) positivity, and Braak score provided by the brain bank of origin. To further dissect the relationships between age, clinical labels of the presence or absence of cognitive impairment, and the slide-level median values of dark blue pixel counts, we used asymptomatic chi square conditional independence tests from the R package bnlearn (v. 4.6.1) [32].

### Code availability

We used the publicly available software tool CLAM [14] to perform deep learning on WSIs and the publicly available software tool HistomicsTK [31] to perform positive pixel counting of the top attention tiles. Scripts used to perform key custom parts of the downstream data analysis are available at the following URL: https://github.com/andymckenzie/deep_histopathology_manuscript.

## Results

### Prediction of cognitive impairment using weakly supervised deep learning

We re-purposed a weakly supervised deep learning algorithm previously used for classification in the setting of a known gold standard label as method for inference of pathophysiology in the setting of noisy cognitive labels (**Fig 1**) [14]. We ran this analysis pipeline on an existing collection of WSIs and trained the model to classify brain tissue sections as coming from the subset of individuals with evidence of antemortem cognitive impairment or not (**Table 1**). In the hippocampus, across the test set of each 10 folds of cross-validation, we found a mean AUC on the 10% of held out test subsets of 0.63 (one-sample Wilcox signed rank test p-value = 0.006; **Fig 2B-C**) and a mean balanced accuracy of 0.59 (p = 0.013, **Fig 2C**). In the frontal cortex, we found a mean AUC of 0.67 (p = 0.002, **Fig 2B-C**) and a mean balanced accuracy of 0.58 (p = 0.009; **Fig 2B**). While the models have modest accuracy as a pure classification task, in both brain regions the classification accuracy was significantly greater than chance, suggesting that the models have utility for the inference of pathophysiology.

**Figure 2.**
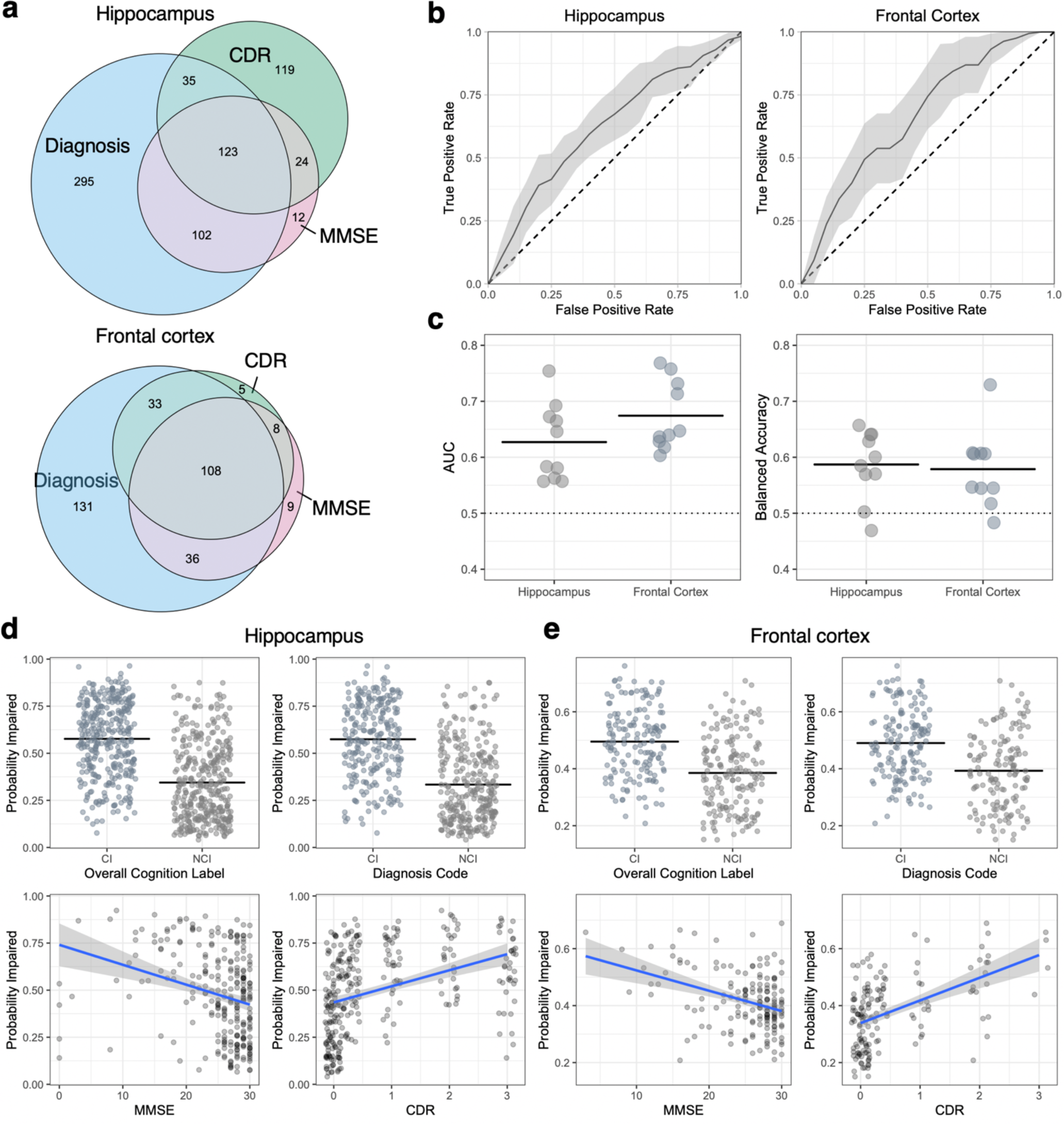
Weakly supervised classification predicts cognitive impairment based on whole slide image data from the hippocampus and frontal cortex. **A**: Venn diagram showing the overlap of the measures used for defining the presence of cognitive impairment in brain donors. **B**: Average receiver operating characteristic curves across 10-fold cross-validation. Error envelopes show ± 1 standard deviation. Horizontal dotted lines show chance-level predictions. **C**: Summary statistics for test evaluation of model performance across 10-fold cross-validation in the frontal cortex and hippocampus. Balanced accuracy refers to the accuracy of predictions weighted by the proportion of labels in both groups in the test split. Horizontal lines are shown at the arithmetic mean values. **D-E**: Probability estimates of cognitive impairment from the top-performing model by each measure of cognitive impairment in the hippocampus (**D**) and frontal cortex (**E**). CDR = Clinical Dementia Rating; MMSE = Mini-Mental State Examination; AUC = Area Under the Curve.

We next evaluated slide-level predictions of the probability of cognitive impairment using the highest performing models in each brain region, parsing out the sub-components of the cognitive impairment classifications (**Fig 2D-E**). In the hippocampus data, we found that the probability of cognitive impairment estimate was significantly associated with the diagnostic category (57% in the CI group vs 33% in the NCI group, t-test p-value < 2.2e-16), MMSE (ρ = -0.32, p = 1.1e-7), and CDR (ρ = 0.5, p < 2.2e-16). In the frontal cortex data, we found that the probability of cognitive impairment estimate was also significantly associated with the diagnostic category (49% in the CI group vs 39% in the NCI group, p = 9.8e-11), MMSE (ρ = -0.30, p = 1.2e-4), and CDR (ρ = 0.52, p = 3.9e-12). While these strong associations with the cognitive labels are expected because they are what the models were trained on, they show that the model has not overly anchored on any one of the three cognitive labels employed. The correlation of the probability estimates of the models from the hippocampus and frontal cortex was highly significant and of moderate strength (ρ = 0.41, p = 1.5e-14; **Supplementary Fig 2**), suggesting that the models trained on the two different brain regions are identifying partially independent signals for cognitive impairment.

To explore the reasons for the imperfect classification accuracy we identified, we found the correlation of the probability estimates of cognitive impairment with age across groups (**Fig 3A-B**). In the hippocampus, there was a significant correlation between age and the estimated probability of cognitive impairment in the non-cognitively impaired group (ρ = 0.37, p = 1.2e-12), a weaker but still significant correlation in the cognitively impaired group (ρ = 0.18, p = 9.0e-4), and a significant difference in correlation (z-score for difference = -2.6; empirical p-value = 0.014). In the frontal cortex, there was a significant correlation between age and the estimated probability of cognitive impairment in the non-cognitively impaired group (ρ = 0.45, p = 1.1e-9), no significant correlation in the cognitively impaired group (ρ = -0.12, p = 0.11), and a significant difference in correlation (z-score = -5.3; empirical p-value = 1e-4). These differential correlation results with age suggests that one reason for imperfect classification accuracy may be mislabeling of brain donors with more advanced age who did have cognitive impairment as not cognitively impaired.

**Figure 3.**
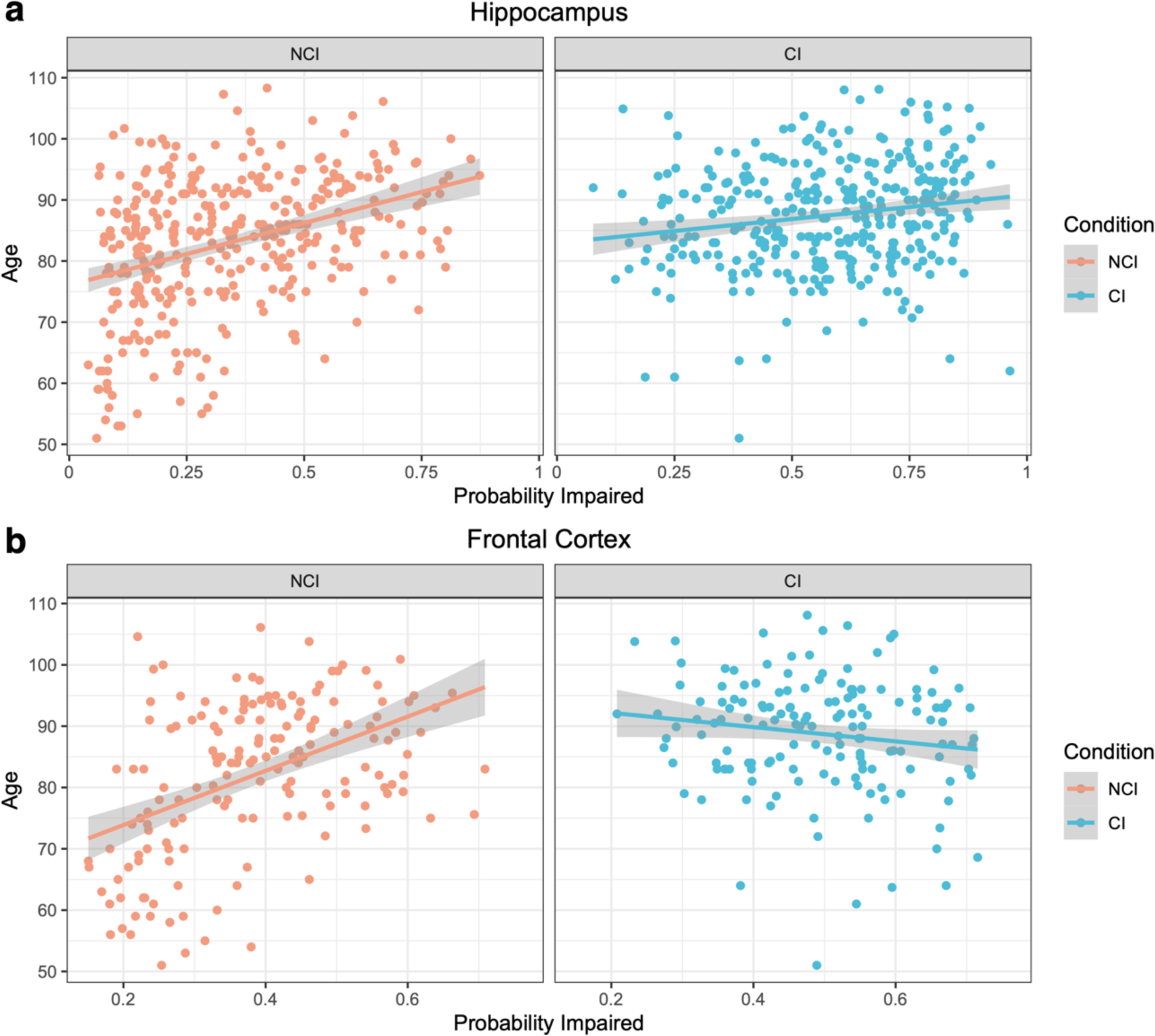
Differential correlations of cognitive impairment probability estimates and age by clinical cognitive impairment label. Scatter plots for the correlation of age and the deep learning model probability estimates for cognitive impairment in the hippocampus (**A**) and frontal cortex (**B**) are shown. Trend lines show predictions using a linear model in each group of data and grey error envelopes show the associated 95% confidence intervals. NCI = Not Cognitively Impaired; CI = Cognitively Impaired.

### Attention-based interpretation identifies an association of white matter pathology with cognitive impairment

To explore the underlying anatomical features used as evidence by the deep learning algorithm, we performed attention-based interpretation analysis using the highest performing models from each brain region. In the hippocampus, on a macro-anatomic scale, the model was found to have qualitatively higher attention in white matter regions as opposed to grey matter (**Fig 4A-B**). On a microanatomic scale, the models were qualitatively found to have a lower level of LFB staining intensity in the hippocampal top attention tiles from the cases labeled with cognitive impairment (**Fig 4C**). To quantify sub-regional differences of the attention signal in the hippocampus, we manually annotated tissue types in a randomly chosen subset of WSIs and used these annotations to measure the region-specific attention scores produced by the model. Quantitative attention scores were found to be significantly higher in the white matter (average attention z-score = 0.62) than in the grey matter (average attention z-score = -0.41; paired t-test for difference p-value = 9.4e-8). The same trend of higher attention scores in the white matter was found across cognitive status labels (**Fig 4D**), suggesting that this result is not due to confounding by cognitive impairment label but instead to properties of the models.

**Figure 4.**
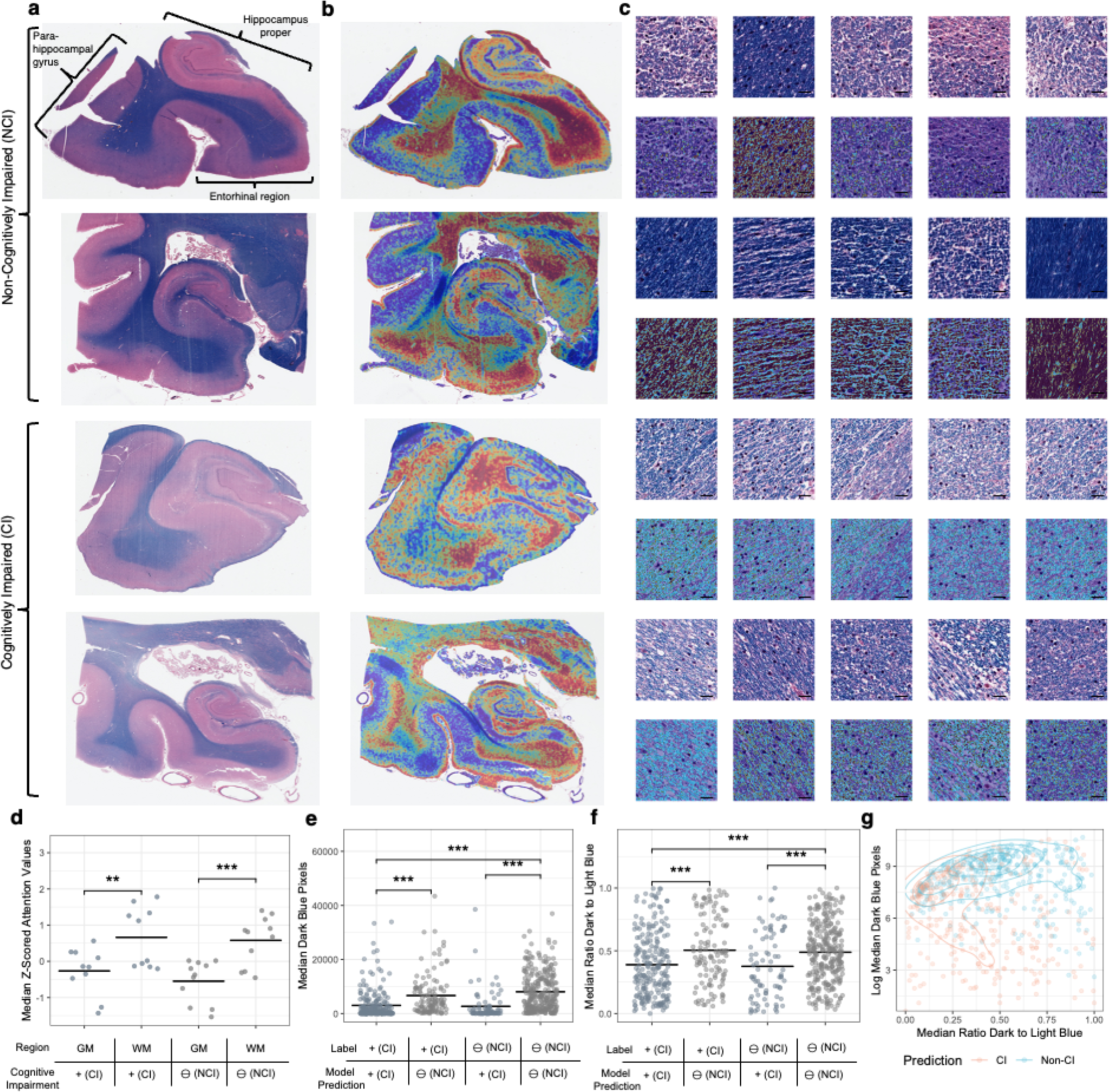
Interpretation of tissue-level attention maps and tile-level staining intensity in the hippocampus suggests myelin loss. **A-B**: Representative WSIs labeled and predicted to be in the non-cognitive impaired (upper) or cognitively impaired (lower) groups (**A**) and corresponding representative attention heatmaps (**B**). In these heatmaps, dark red indicates high attention, while dark blue indicates low attention. **C**: Top 5 highest attention tiles (upper) and blue hue range positive pixel annotations (lower) from the matching WSIs as shown in sub-figures A/B. Scale bar = 20 μm. **D**: Median z-transformed attention score values in the grey matter and white matter. Each data point is a median attention score from the white matter or the grey matter from one WSI. **E-F**: Median dark blue range pixel counts as a measure of LFB staining intensity (**E**) and ratio of the dark blue to light blue pixel counts in the top attention tiles of WSIs predicted and labeled to have cognitive impairment or not (**F**). **G**: Scatter plot and contour lines showing the relationship between dark blue range pixel counts and the ratio of the dark blue to light blue pixel counts in the top attention tiles of WSIs. Orange dots indicate that the WSI was predicted to come from a CI donor, while blue dots indicate NCI. * = p < 0.05, *** = p < 0.001. GM = Grey Matter; WM = White Matter; CI = Cognitively Impaired; NCI = Not Cognitively Impaired.

To quantify the microanatomic scale results from the hippocampus, we used positive pixel counting on the 100 tiles with the highest attention scores as measured by the model, henceforth called the “top tiles.” LFB stains CNS myelin sheaths dark blue [24] and our chosen pixel range was designed to capture this LFB staining intensity (**Supplementary Fig 1**). In the hippocampus, the WSIs predicted to be from brain donors with cognitive impairment had a significantly lower LFB staining intensity in the top tiles (t-test difference p-value = 7.6e-7, **Fig 4E**). To normalize for possible variation in staining intensity across slides, we measured the ratio of dark blue staining to light blue staining in the top 100 attention tiles. We found that there was a significantly lower ratio of dark blue staining to light blue staining (t-test p = 4.1e-5; **Fig 4F**). The LFB staining intensity and the ratio of dark blue to light blue staining intensity in the top tiles are correlated (ρ = 0.14, p = 2.8e-4) and jointly distinguish donors predicted by the model to be cognitively impaired or not (**Fig 4G**).

We next performed the same analysis in the frontal cortex data set, where the results largely echoed those of the hippocampus, with generally stronger effect sizes. Qualitatively, the frontal cortex model was also found to have higher attention in white matter regions (**Fig 5A-B**) and a lower level of LFB staining in the top tiles (**Fig 5C**). Quantitatively, attention scores were found to be significantly higher in the white matter (average attention z-score =1.03) than in the grey matter (median attention z-score = -0.85, t-test p-value < 2.2e-16; **Fig 5D**). The group labeled as cognitively impaired had a significantly lower LFB intensity in the top tiles (t-test p = 7.3e-6, **Fig 5E**) and there was a significantly lower ratio of dark blue staining to light blue staining (t-test p < 2.2e-16, **Fig 5F**). And as with the hippocampus data set, these two measures are correlated and jointly distinguish between brain donors with and without labels of cognitive impairment (**Fig 5G**).

**Figure 5.**
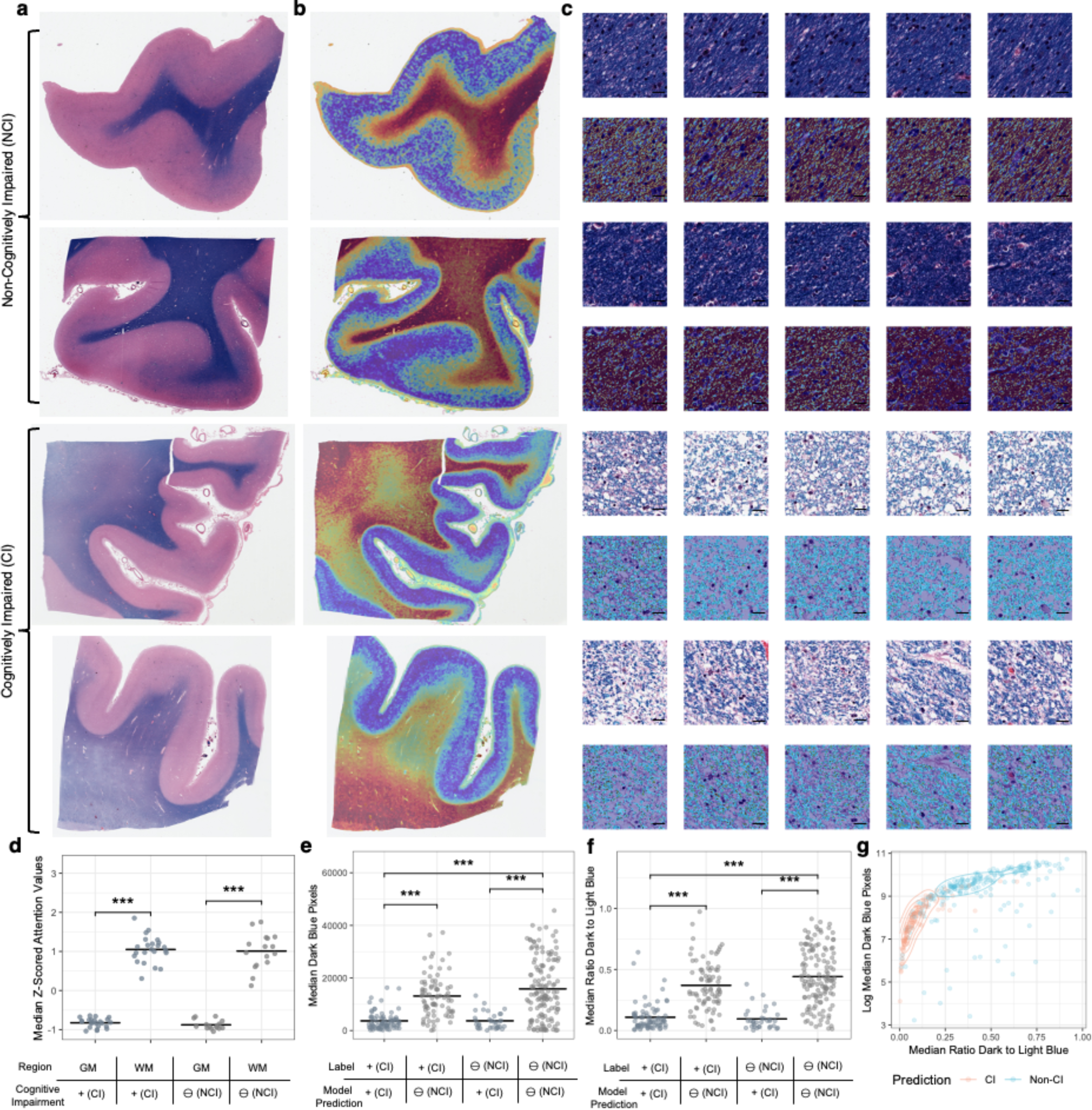
Interpretation of attention maps and tile-level myelin density in the frontal cortex suggests myelin loss. **A-B**: Representative WSIs labeled and predicted to be in the non-cognitive impaired (upper) or cognitively impaired (lower) groups (**A**) and corresponding representative attention heatmaps (**B**). In these heatmaps, dark red indicates high attention while dark blue indicates low attention. **C**: Top 5 highest attention tiles (upper) and blue hue range positive pixel annotations (lower) from the matching WSIs as shown in sub-figures A/B. Scale bar = 20 μm. **D**: Median z-transformed attention score values in the grey matter and white matter. Each data point is a median attention score from the white matter or the grey matter from one WSI. **E-F**: Median dark blue range pixel counts as a measure of LFB staining intensity (**E**) and ratio of the dark blue to light blue pixel counts in the top attention tiles of WSIs predicted and labeled to have cognitive impairment or not (**F**). **G**: Scatter plot and contour lines showing the relationship between dark blue range pixel counts and the ratio of the dark blue to light blue pixel counts in the top attention tiles of WSIs. Orange dots indicate that the WSI was predicted to come from a CI individual, while blue dots indicate NCI. *** = p < 0.001. GM = Grey Matter; WM = White Matter; CI = Cognitively Impaired; NCI = Not Cognitively Impaired.

### Deep histopathological findings are partially independent of several known pathoanatomic features

We compared the deep learning model results with previously established clinicopathologic features, namely age, Braak stage, cerebrovascular pathology, and hippocampal ARTAG. This association analysis was focused on the hippocampal data set because it has a substantially higher sample size and is therefore better powered to detect correlations (**Fig 6A**). We found that there was a significant rank correlation of the model’s cognitive impairment probability estimates with age (ρ = 0.32, p < 2.2e-16), Braak stage (ρ = 0.13, p = 6.2e-4), ARTAG positivity (ρ = 0.15, p = 1.5e-4), and cerebrovascular pathology (ρ = 0.29, p = 9.0e-5). We also found that there was a significant association of LFB staining intensity in the top attention tiles with age (ρ = -0.18, p = 1.0e-6) and cerebrovascular pathology (ρ = -0.24, p = 0.0014), but not with Braak stage (ρ = -0.05, p = 0.18) or with the presence of ARTAG pathology (ρ = -0.05, p = 0.24). This result suggests that the deep learning algorithm has identified a signal for cognitive impairment that is associated with some aspects of known pathophysiology.

**Figure 6.**
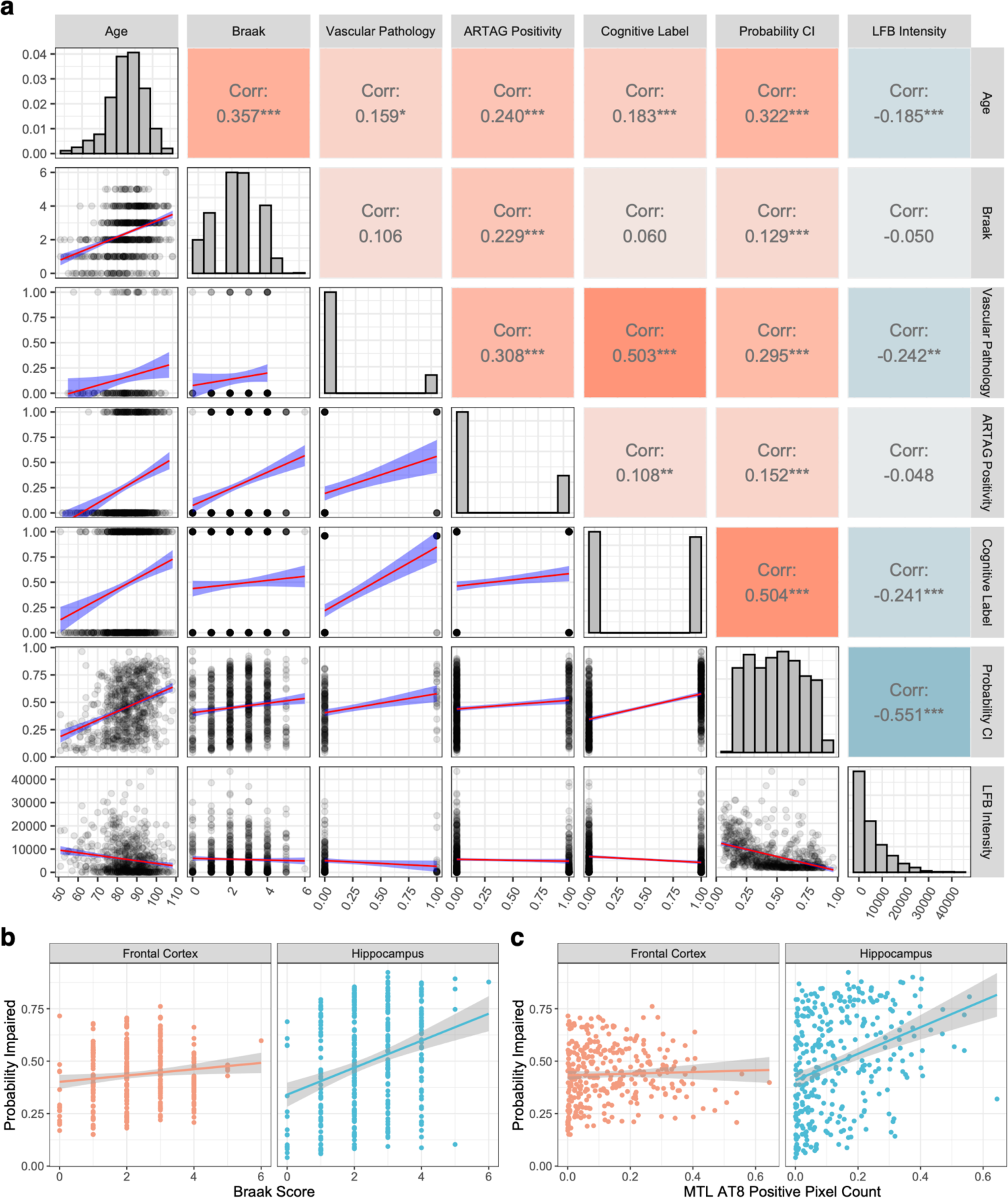
Deep histopathology features are partially associated with several known clinicopathologic features and partially independent. **A**: Correlation analysis of deep histopathology results and clinicopathologic features: age, Braak score, evidence of cerebrovascular pathology (coded as 0 = not present and 1 = present), ARTAG positivity in the hippocampus (coded as 0 = not present and 1 = present), cognitive label (coded as 0 = not cognitively impaired and 1 = cognitively impaired), probability of cognitive impairment as predicted by the top-performing model trained on the hippocampal data, and median LFB staining intensity in the top attention tiles in the hippocampus data set. Upper right: rank correlation values and associated p-values (* = p < 0.05, ** = p < 0.01, *** = p < 0.001). Diagonal: histograms of variables. Lower left: scatterplots with linear model trend lines for the variable pairs (red lines) and 95% confidence intervals (blue envelopes). This plot was made using the R package GGally (v. 2.1.2). **B-C**: Scatter plots for probability of cognitive impairment estimated in the frontal cortex and hippocampus with Braak stage (**B**) and AT8 staining positive pixel counts in the medial temporal lobe (MTL) (**C**). Trend lines show predictions via a linear model and grey envelopes show associated 95% confidence intervals. CI = Cognitive impairment; ARTAG = Aging-related tau astrogliopathy; LFB = Luxol Fast Blue.

We further dissected the association between age, LFB staining intensity in the top tiles, and cognitive impairment labels in the hippocampal data set with conditional independence tests. We found that the label of cognitive impairment was not conditionally independent of age when accounting for LFB staining intensity in the top tiles (mutual information = 42.3, p = 3.4e-9 by asymptomatic chi square test). Additionally, the label of cognitive impairment was not conditionally independent of LFB staining intensity in the top tiles when accounting for age (mutual information = 21.8, p = 7.1e-5). These results suggest that while these three variables are all significantly associated with one another, chronological age does not fully explain the association of LFB staining intensity in the top tiles with cognitive impairment, nor vice versa.

We next performed correlation analysis on the frontal cortex data set (**Supplementary Fig 3**), omitting cerebrovascular pathology as a variable because the intersected sample size was too low for reliable estimates in the frontal cortex data set. While the results between brain regions were predominantly similar, one difference is that there was not a significant correlation identified between the model’s cognitive impairment probability estimates and Braak stage in the frontal cortex, although it trended towards significance (ρ = 0.11, p = 0.06). Because the hippocampus has a larger sample size than the frontal cortex, it is better powered to detect a significant correlation between Braak stage and probability of cognitive impairment. To address the possibility that this difference in sample size affected any differences in correlation between the regions, we filtered the sample to select only those cases containing data from both the hippocampus and frontal cortex and tested for a differential correlation. In this subset of the data, we found a higher rank correlation between Braak stage and the probability of cognitive impairment derived from the hippocampus (ρ = 0.29, p = 7.4e-8) than in the frontal cortex (ρ = 0.11, p = 0.06), which was a significant difference in correlation (z-score for difference = -2.4, empirical p-value = 0.02; **Fig 6B**). In order to query the robustness of this result, we employed data on positive pixel counts for AT8 staining in the medial temporal lobe (MTL), a measure of tau burden that has been previously described in this cohort [22]. We found that there was a significant rank correlation between AT8 staining burden in the MTL and the probability of cognitive impairment derived from the hippocampus (ρ = 0.37, p = 9.2e-12), a weaker but still significant correlation with the probability of cognitive impairment derived from the frontal cortex (rho = 0.12, p = 0.029), and that there was a significantly higher correlation between these two measures in the hippocampus (z-score for difference = 3.2, empirical p-value = 0.001; **Fig 6C**). One way to interpret these findings is that the contributions of different types of histopathology to the deep learning-derived predicted probability of cognitive impairment may differ by brain region.

## Discussion

In this study, we used deep learning models to identify a reduction in LFB staining intensity in the top attention tiles from brain sections of donors with antemortem evidence of cognitive impairment. Because LFB staining in brain tissue is generally used to quantify the amount of myelin [24, 25], the signal that we identified is likely due to decreased myelin staining intensity. Our results are not able to distinguish decreased myelin density with spared axons as opposed to axon injury and associated myelin loss. In many cases, diminished myelin density in aging is associated with cerebrovascular disease [33]. This is consistent with the strong correlations we identified in this study between cerebrovascular pathology, the predicted probability of cognitive impairment, and decreased LFB staining in the top attention tiles. Even when accounting for age, there was still an association between decreased LFB staining in the top attention tiles and cognitive impairment. Treating cerebrovascular disease risk factors such as hypertension has been found to decrease white matter pathology and partially reverse age-related cognitive impairment [33]. However, age-associated decreases in myelin density have numerous possible causes other than cerebrovascular disease, such as nearby AD cortical pathology [34, 35], a primary effect of aging [36–38], repetitive head impacts [39], or the accumulated effects of excessive alcohol use [40]. It is unclear the extent to which the decreased myelin density we found to be associated with age-related cognitive impairment are explained solely by cerebrovascular pathology as opposed these other possible etiologies, which warrants further investigation.

While postmortem brain gene expression and pathoanatomical studies in aging and AD have often focused on grey matter, neuroimaging findings over the past several decades have frequently found alterations in the white matter to be strongly associated with cognitive impairment [41]. Leukoaraiosis (*leuko* – white, *araiosis* – rarefaction) is a common neuroimaging abnormality of the white matter that can be found in periventricular or subcortical areas [42, 43]. On T2-weighted and FLAIR MRI, leukoaraiosis is frequently described as white matter hyperintensities [44]. While leukoaraiosis is strongly associated with cerebrovascular disease, the precise etiology remains unclear [43, 44]. Clinically, leukoaraiosis is associated with cognitive deficits such as bradyphrenia [33]. Histologically, leukoaraiosis has been suggested to be associated with decreased density of myelin sheaths [45]. Our deep learning models identified a neurohistologic signal for cognitive impairment that was (a) focused in the white matter, (b) in some cases scattered in a non-uniform pattern across the tissue, (c) and associated with decreased myelin staining intensity. Although our data set lacks associated *in vivo* neuroimaging data to draw conclusive statements, one clear possibility is that the white matter histologic alterations the deep learning models identified may reflect similar etiopathology as the neuroimaging finding of leukoaraiosis. We propose that diminished LFB staining intensity in particular areas identified by a deep learning model may be a quantitative way to assess for the presence of leukoaraiosis-associated neuropathology in postmortem brains.

It is important to consider the limitations of this study. First, compared to previously published weakly supervised learning publications in oncology (which are often n > 1000), the data set employed here (*n* = 716) is not as large [13, 14]. Because there is an absence of significant Aβ burden in this cohort, it also limits the representativeness of the cohort to the population at large. This adds to the numerous selection biases in brain donation-based autopsy cohorts in general [46]. Second, the WSI data set analyzed only contains one stain, the LH&E stain. While LFB staining is ideal for detecting myelin, it is possible that it may have highlighted the white matter to a disproportionate degree that affected the deep learning algorithm results. Third, we were unable to assess for comorbid TDP-43 pathology, which would allow us to screen for limbic-predominant age-related TDP-43 encephalopathy (LATE), a common TDP-43 proteinopathy associated with an amnestic dementia in elderly individuals [47]. Additionally, because we only looked at two brain regions, we have limited anatomical sampling, which is problematic because we know that cognitive impairment is determined by accumulated lesion burden across the brain. Finally, it is possible that a fixation or staining artifact may affect the cognitive impairment probability estimates and/or attention signals. For example, deeper areas of the brain were often found to have qualitatively higher attention signals, which may have been artifactual. However, this is considered less likely as – again qualitatively – the attention signal appears to follow anatomical compartments, such as the deep white matter, while sparing the subcortical U-fibers, regardless of the depth of these compartments.

Although there are some additional limitations to our study, we expect that our methodology lays the groundwork for further probing of the histopathology of age-related cognitive impairment in future studies that will be able to address these limitations. For example, while our current slide-level predictive accuracy is modest, as our data sets grow, we expect it will improve in discriminative power and will better enable us to pinpoint morphological features associated with cognitive impairment. Additionally, while we only focused on a robust yet general approach to assessing myelin, *i.e.* LH&E stained tissue sections, future studies deploying additional modalities of assessing myelin injury, such as immunohistochemical staining for oligodendrocyte, axonal, vascular, inflammatory, and myelin markers, will help to further elucidate the pathogenesis of age-related cognitive impairment. Finally, because we only have WSI data available from two brain regions, we are limited in our ability to explain why the results from the two brain regions appeared to differ in some ways. Our ability to interpret differences in deep learning-derived metrics across brain regions will improve with richer data sets containing WSIs from more brain regions.

As compared to cancer pathology, which has heretofore been the main use case of weakly supervised deep learning in digital pathology, in studying the neuropathology of dementia, there is less of an emphasis on diagnosis and more of an emphasis on the inference of pathophysiology. This is in part because cancer can be more frequently associated with one causal type, whereas cognitive deficits in the brain are generally due to overlapping pathologies with complex patterns of comorbidity. The multifactorial nature of cognitive deficits lends itself well to multidimensional interpretation studies. First, it emphasizes the value of quantitative probability estimates of cognitive impairment instead of binary labels, which allow for more precise correlation analysis with other clinicopathologic features. Second, the relative focus on understanding pathophysiology in neuropathology also underscores the value of deterministic computer vision studies, such as positive pixel counting, as a downstream method for interrogating attention or other interpretability signals present in deep learning models. While the prediction capacity of deep learning models in digital pathology can be expected to continue to improve rapidly, our ability to understand what histopathologic features those models are focused on is lagging.

Improving our suite of methods for the interpretation of deep learning models will allow us to best harness them and to understand how they may be flawed or biased. Because the study of the neuropathology of dementia remains driven by human ingenuity, more interpretable deep learning methods will be essential to accelerate its adoption across the field.

Predicting the presence or absence of cognitive impairment with the use of single histology sections on an individual level is an extremely challenging task. There are known barriers related to disease heterogeneity, variation in clinician practices, and cognitive reserve [8, 48]. In this study, we employed a deep learning classification model for inference of pathophysiology from histology slides with noisy labels of cognitive impairment, resulting in predictions with modest accuracy but significantly above chance level. Interpretation studies suggested that top performing models in the hippocampus and frontal cortex focused on similar aspects of white matter pathology. On a macroanatomic level, they had higher attention on white matter than gray matter; on a microanatomic level, the highest attention tiles showed differences in LFB staining intensity between slides from brains donors predicted to have cognitive impairment or not. Both the probability estimates of cognitive impairment and the measure of LFB staining intensity in the top attention tiles were partially independent of several known pathoclinical features, suggesting that they may be identifying unexpected aspects of pathophysiology. On the other hand, the probability estimates of cognitive impairment were not completely explained by LFB intensity in the top attention tiles; for example, ARTAG positivity was significantly associated with the probability estimates of cognitive impairment from the deep learning models but not with LFB intensity in the top attention tiles. Our results demonstrate that weakly supervised deep learning is a promising approach to dissect pathoanatomic features associated with cognitive deficits in neurohistologic data sets in an unbiased manner.

## Acknowledgments

This work was supported in part through the computational resources and staff expertise provided by Scientific Computing at the Icahn School of Medicine at Mount Sinai. Additionally, this work was supported by an Alzheimer’s Disease Research Center (ADRC) Developmental Project Funding Award to A.T.M (P30 AG066514).

*The PART working group is: Jean-Paul Vonsattel, Andy F. Teich (Columbia University); Marla Gearing, Jonathan Glass (Emory University); Juan C. Troncoso (Johns Hopkins University), Matthew P. Frosch, Bradley T. Hyman (Harvard Medical School and Massachusetts General Hospital, the Massachusetts Alzheimer Disease Research Center); Dennis W. Dickson, Melissa E. Murray (Mayo Clinc), Johannes Attems (Newcastle Brain Tissue Resource, Newcastle University); Margaret E. Flanagan, Qinwen Mao, M-Marsel Mesulam, Sandra Weintraub (Northwestern University); Randy L. Woltjer, Thao Pham (Oregon Health Sciences University); Julia Kofler (University of Pittsburgh), Julie A. Schneider, Lei Yu (Rush University); Dushyant P. Purohit, Vahram Haroutunian, Patrick R. Hof, Sam Gandy, Mary Sano (Icahn School of Medicine at Mount Sinai); Thomas G. Beach (Banner Sun Health); Wayne Poon, Claudia Kawas, María Corrada (The Brain Repository at the University of California Irvine); Robert A. Rissman, Jeff Metcalf, Sara Shuldberg, Bahar Salehi (University of California San Diego); Peter T. Nelson (University of Kentucky); John Q. Trojanowski, Edward B. Lee, David A Wolk, Corey T McMillan (University of Pennsylvania); C. Dirk Keene, Caitlin S. Latimer, Thomas J. Montine (The BioRepository and Integrated Neuropathology Laboratory and Precision Neuropathology Core at the University of Washington); Gabor G. Kovacs, Mirjam I. Lutz, Peter Fischer (Medical University of Vienna); Richard J. Perrin, Nigel J. Cairns, Erin E. Franklin (Knight Alzheimer Disease Research Center Neuropathology Core at Washington University School of Medicine). The authors would also like to acknowledge Ping Shang, HT(ASCP) QIHC and Jeff Harris, HTL(ASCP) for histologic and immunohistochemical preparations, and Chan Foong, M.S., for preparation of whole slide images (University of Texas Southwestern), as well as the Neuropathology Brain Bank & Research Core at Mount Sinai.

Grants: MSSM: Alzheimer’s Disease Research Center at Mount Sinai Developmental Project Funding Award (ATM), R01 AG054008, R01 NS095252, R01 AG060961, and R01 NS086736 Rainwater Charitable Foundation, Alexander Saint-Amand Fellowship (JFC), F32 AG056098 and P30 AG066514 (KF), P50 AG005138 and P30 AG066514 (VH, JFC, MS, SG, AG, PH), 75N95019C00049 (VH), BU / MSSM / MAYO: R01 AG062348 (AM JFC DD) CUMC: P50AG008702 (JPV AFT) BU: U54 NS115266 (AM) R01 CA079830 (HTC) UPENN: P30 AG010124, P01 AG017586 and U19 AG062418 (JQT), P30 AG072979 and P01 AG066597 (EBL), R01 AG066152 (CTM) PITT: R01 AG066152 P30 AG066468 (JK) Banner: U24 NS072026 and P30 AG019610 The Arizona Department of Health Services, and the Michael J. Fox Foundation for Parkinson’s Research (TB) Northwestern: P30 AG013854 (MEF) Emory: P30 NS055077 and P50 AG025688 (MG) OHSU: P30 AG08017 (RW) UTSW: Winspear Family Center for Research on the Neuropathology of Alzheimer Disease (CWIII) MADRC: BT P50 AG05134 (BH) RUSH: ADC grant AG10161 and MAP grant (JS) UCI: R01AG021055 and P50AG016573 (CK) P01AG000538 (WP) UCSD: P30 AG062429 01 and P50 AG005131 (RR) UK: P30 AG028383 (PN) U Washington: P50 AG005136, P30 AG066509, U01 AG006781, U19 066567 and the Nancy and Buster Alvord Endowment (CKD) Washington U / Knight ADRC: P30 AG066444, P01 AG003991 and P01 AG026276 (CDK) Newcastle: UK Medical Research Council (G0400074), by Brains for Dementia research, a joint venture between Alzheimer’s Society and Alzheimer’s Research UK and by the NIHR Newcastle Biomedical Research Centre awarded to the Newcastle upon Tyne Hospitals NHS Foundation Trust and Newcastle University (JA). A portion of the human tissue used in the study was obtained from the NIH NeuroBioBank.

## Author Contributions

A.T.M., G.M., K.F., and J.F.C. contributed to the original design of the study; J.M.W., T.E.R., and C.L.W. contributed tissue processing; A.T.M., G.M., K.F., and D.K. contributed coding; K.F., M.A.I., and J.F.C. contributed to pathoclinical data collection for the cohort; G.M., M.S., and J.F.C. contributed annotations; G.C. and T.J.F. contributed deep learning data analysis; J.M.W., T.E.R., A.C.M., T.D.S., and C.L.W. contributed neuropathology data analysis; A.T.M., G.C., K.F., and J.F.C. wrote the original draft of the manuscript. All authors read and approved the manuscript.

## Data availability

All whole slide image and matched pathoclinical data is available to the academic community upon request.

## Supplementary Figures

**Supplementary Figure 1.**
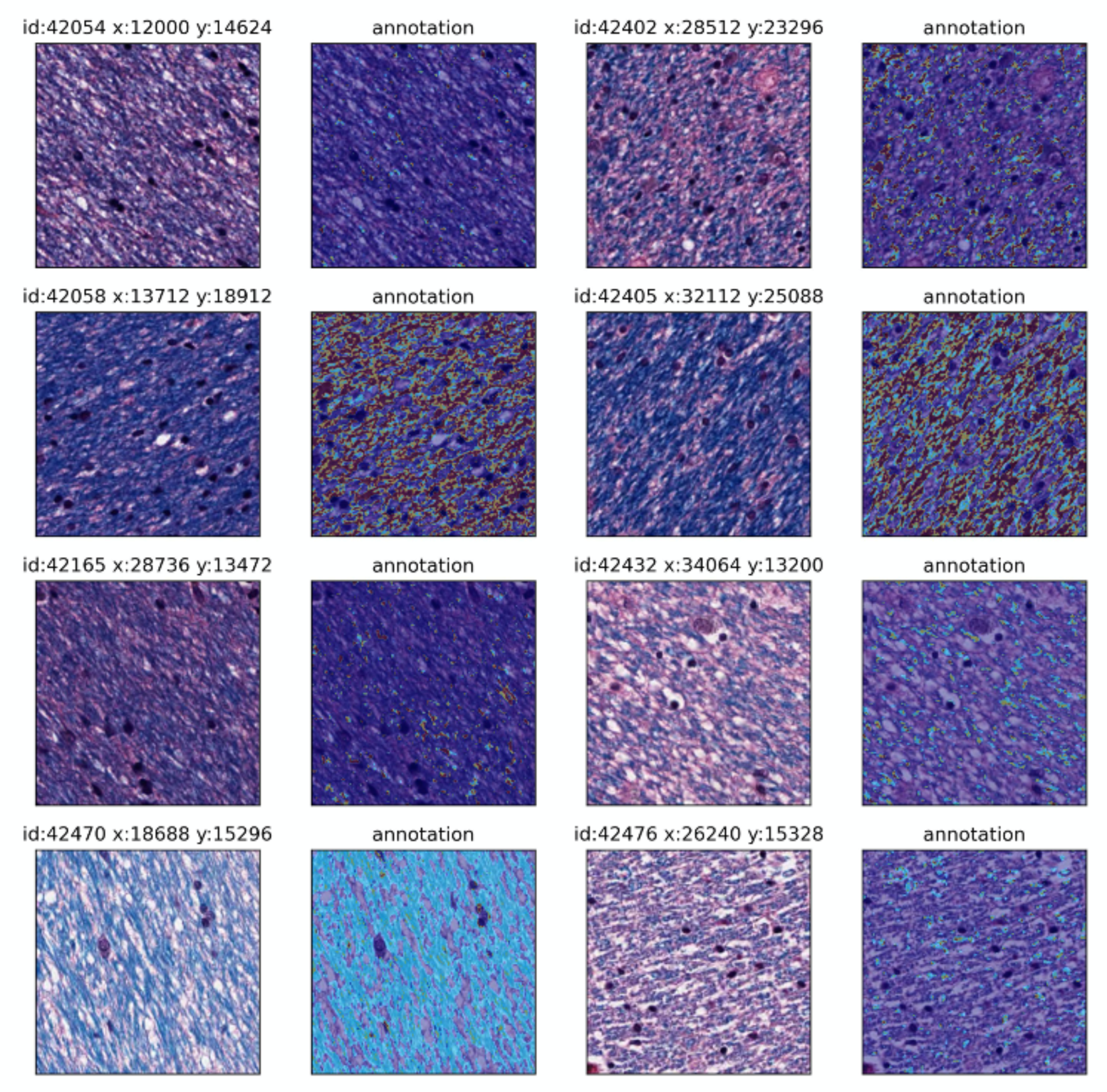
Annotation procedure of blue hue ranges in Luxol fast blue, hematoxylin, and eosin-stained tiles. Representative tiles show the annotation method used for positive pixel counting in the Luxol fast blue, hematoxylin, and eosin (LH&E) stained histology tiles. For the annotation heatmap, the darker blue pixel range is highlighted as red while the lighter blue pixel range is highlighted as light blue.

**Supplementary Figure 2.**
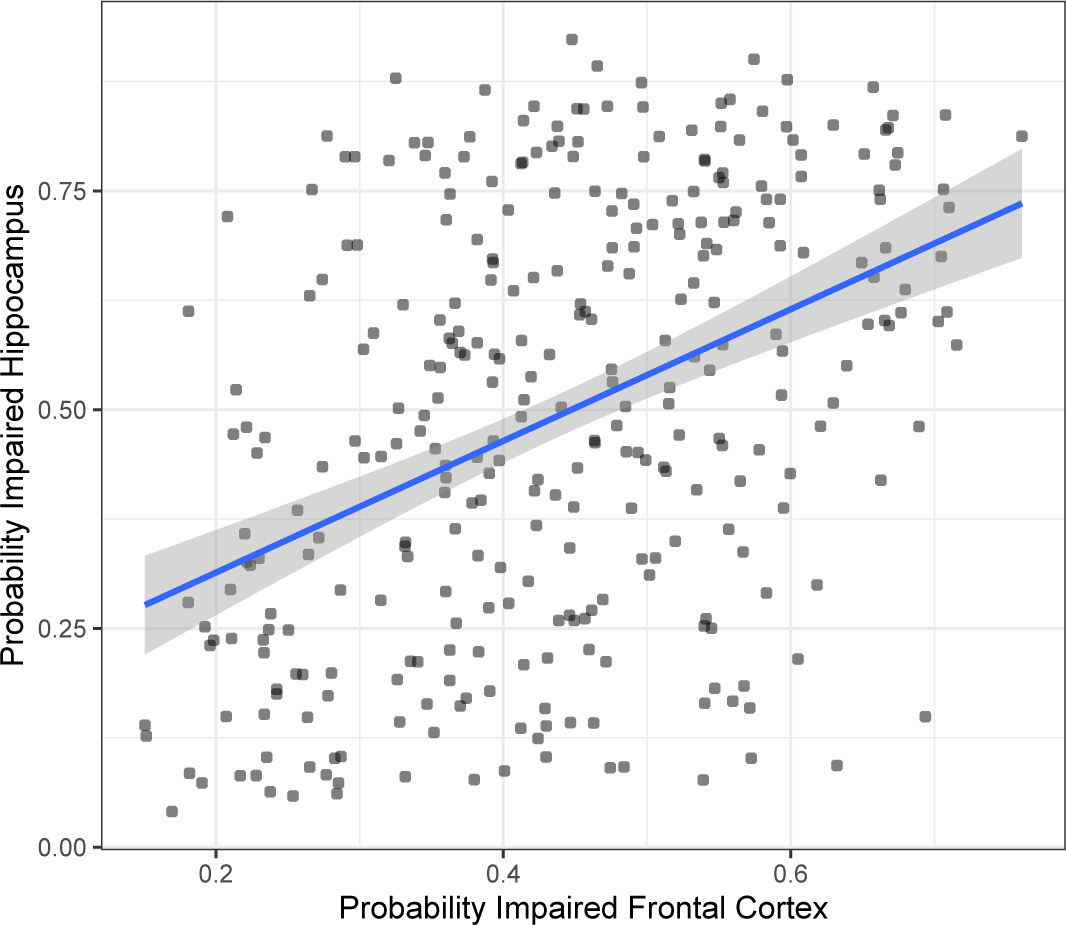
Correlation of slide-level probability estimates of cognitive impairment in matched brain donors between the two brain regions. Scatter plots showing the probability estimates of cognitive impairment by the top-performing models in the same brain donors between WSIs in the hippocampus and frontal cortex data sets. The blue line shows predictions from a linear model and grey error envelopes show 95% confidence intervals for the linear model.

**Supplementary Figure 3.**
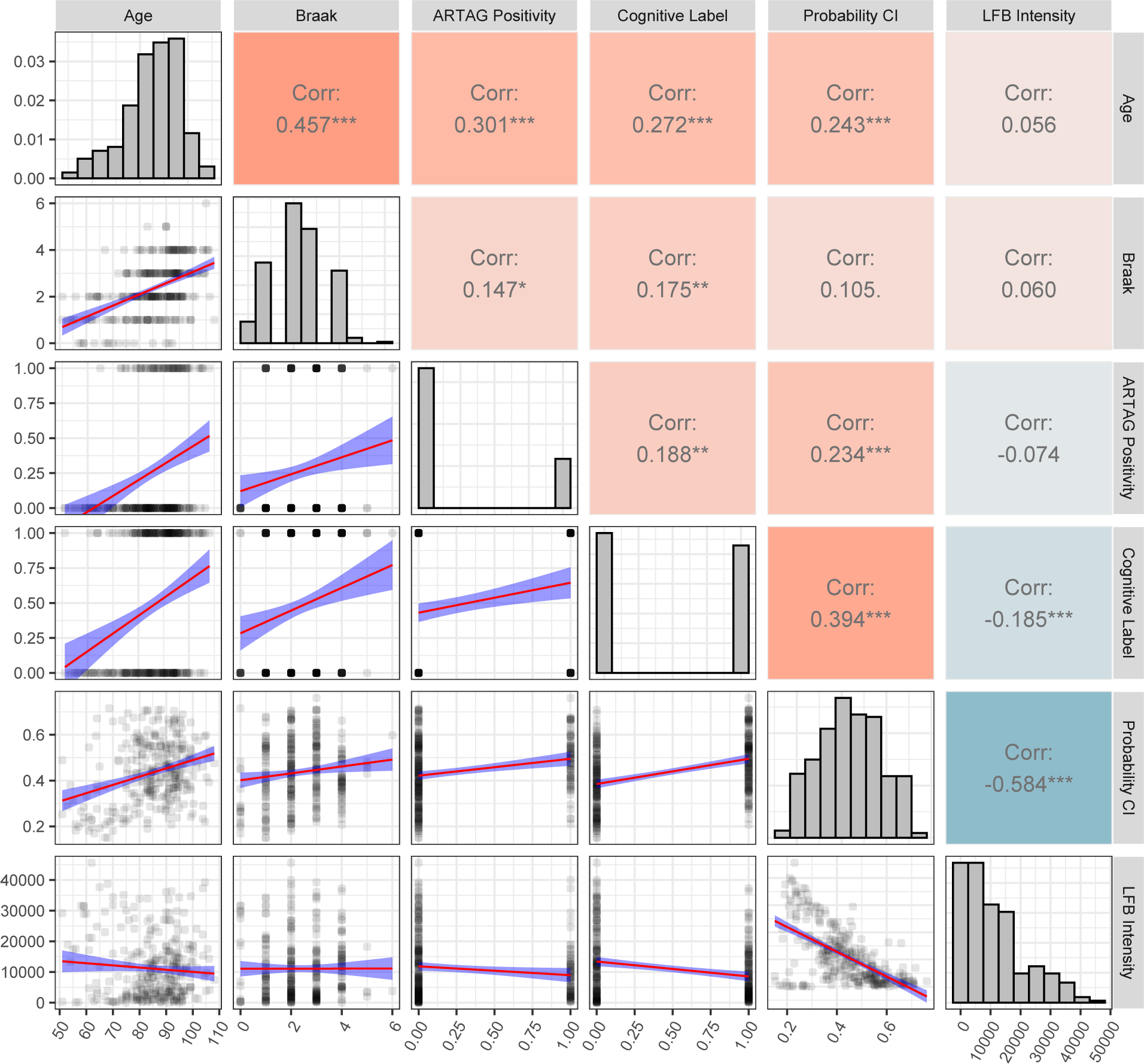
Scatterplot matrix of deep histopathology features with clinicopathologic features in the frontal cortex. Correlation analysis of deep histopathology results and clinicopathologic features: age, Braak score, ARTAG positivity in the hippocampus (coded as 0 = not present and 1 = present), cognitive label (coded as 0 = not cognitively impaired and 1 = cognitively impaired), probability of cognitive impairment as predicted by the top-performing model trained on the frontal cortex data, and median LFB staining intensity (pixel counts) in the top attention tiles in the frontal cortex data set. Upper right: rank correlation values and associated p-values (* = p < 0.05, ** = p < 0.01, *** = p < 0.001). Diagonal: histograms of variables. Lower left: Scatterplots with linear model trend lines for the variable pairs (red lines) and 95% confidence intervals (blue envelopes). This plot was made using the R package GGally (v. 2.1.2). CI = Cognitive impairment; ARTAG = Aging-related tau astrogliopathy; LFB = Luxol Fast Blue.

## References

1. Kaup AR, Mirzakhanian H, Jeste DV, Eyler LT. A Review of the Brain Structure Correlates of Successful Cognitive Aging. J Neuropsychiatry Clin Neurosci. 2011;23:6–15.

2. Ackley SF, Zimmerman SC, Brenowitz WD, Tchetgen EJT, Gold AL, Manly JJ, et al. Effect of reductions in amyloid levels on cognitive change in randomized trials: instrumental variable meta-analysis. BMJ. British Medical Journal Publishing Group; 2021;372:n156.

3. Matthews FE, Brayne C, Lowe J, McKeith I, Wharton SB, Ince P. Epidemiological pathology of dementia: attributable-risks at death in the Medical Research Council Cognitive Function and Ageing Study. PLoS Med. 2009;6:e1000180.

4. Schneider JA, Arvanitakis Z, Bang W, Bennett DA. Mixed brain pathologies account for most dementia cases in community-dwelling older persons. Neurology. 2007;69:2197–204.

5. Kapasi A, DeCarli C, Schneider JA. Impact of multiple pathologies on the threshold for clinically overt dementia. Acta Neuropathol (Berl). 2017;134:171–86.

6. Power MC, Mormino E, Soldan A, James BD, Yu L, Armstrong NM, et al. Combined neuropathological pathways account for age-related risk of dementia. Ann Neurol. 2018;84:10–22.

7. Murman DL. The Impact of Age on Cognition. Semin Hear. 2015;36:111–21.

8. Nelson PT, Alafuzoff I, Bigio EH, Bouras C, Braak H, Cairns NJ, et al. Correlation of Alzheimer disease neuropathologic changes with cognitive status: a review of the literature. J Neuropathol Exp Neurol. 2012;71:362–81.

9. Aeffner F, Zarella MD, Buchbinder N, Bui MM, Goodman MR, Hartman DJ, et al. Introduction to Digital Image Analysis in Whole-slide Imaging: A White Paper from the Digital Pathology Association. J Pathol Inform. 2019;10:9.

10. Signaevsky M, Prastawa M, Farrell K, Tabish N, Baldwin E, Han N, et al. Artificial intelligence in neuropathology: deep learning-based assessment of tauopathy. Lab Invest. 2019;99:1019–29.

11. Tang Z, Chuang KV, DeCarli C, Jin L-W, Beckett L, Keiser MJ, et al. Interpretable classification of Alzheimer’s disease pathologies with a convolutional neural network pipeline. Nat Commun. 2019;10:2173.

12. Vega AR, Chkheidze R, Jarmale V, Shang P, Foong C, Diamond MI, et al. Deep learning reveals disease-specific signatures of white matter pathology in tauopathies. Acta Neuropathol Commun. 2021;9:170.

13. Campanella G, Hanna MG, Geneslaw L, Miraflor A, Werneck Krauss Silva V, Busam KJ, et al. Clinical-grade computational pathology using weakly supervised deep learning on whole slide images. Nat Med. 2019;25:1301–9.

14. Lu MY, Williamson DFK, Chen TY, Chen RJ, Barbieri M, Mahmood F. Data-efficient and weakly supervised computational pathology on whole-slide images. Nat Biomed Eng. 2021;

15. Bell V, Wilkinson S, Greco M, Hendrie C, Mills B, Deeley Q. What is the functional/organic distinction actually doing in psychiatry and neurology? Wellcome Open Res. 2020;5:138.

16. DeTure MA, Dickson DW. The neuropathological diagnosis of Alzheimer’s disease. Mol Neurodegener. 2019;14:32.

17. Rabinovici GD, Carrillo MC, Forman M, DeSanti S, Miller DS, Kozauer N, et al. Multiple comorbid neuropathologies in the setting of Alzheimer’s disease neuropathology and implications for drug development. Alzheimers Dement N Y N. 2017;3:83–91.

18. Houx PJ, Shepherd J, Blauw G-J, Murphy MB, Ford I, Bollen EL, et al. Testing cognitive function in elderly populations: the PROSPER study. PROspective Study of Pravastatin in the Elderly at Risk. J Neurol Neurosurg Psychiatry. 2002;73:385–9.

19. Lim ASP, Gaiteri C, Yu L, Sohail S, Swardfager W, Tasaki S, et al. Seasonal plasticity of cognition and related biological measures in adults with and without Alzheimer disease: Analysis of multiple cohorts. PLoS Med. 2018;15:e1002647.

20. Patnode CD, Perdue LA, Rossom RC, Rushkin MC, Redmond N, Thomas RG, et al. Screening for Cognitive Impairment in Older Adults: Updated Evidence Report and Systematic Review for the US Preventive Services Task Force. JAMA. 2020;323:764–85.

21. Farrell K, Kim S, Han N, Iida MA, Gonzalez EM, Otero-Garcia M, et al. Genome-wide association study and functional validation implicates JADE1 in tauopathy. Acta Neuropathol (Berl). 2022;143:33– 53.

22. Iida MA, Farrell K, Walker JM, Richardson TE, Marx GA, Bryce CH, et al. Predictors of cognitive impairment in primary age-related tauopathy: an autopsy study. Acta Neuropathol Commun. 2021;9:134.

23. Walker JM, Richardson TE, Farrell K, Iida MA, Foong C, Shang P, et al. Early Selective Vulnerability of the CA2 Hippocampal Subfield in Primary Age-Related Tauopathy. J Neuropathol Exp Neurol. 2021;80:102–11.

24. Carriel V, Campos A, Alaminos M, Raimondo S, Geuna S. Staining Methods for Normal and Regenerative Myelin in the Nervous System. Methods Mol Biol Clifton NJ. 2017;1560:207–18.

25. Scholtz CL. Quantitative histochemistry of myelin using Luxol Fast Blue MBS. Histochem J. 1977;9:759–65.

26. McKee AC, Stern RA, Nowinski CJ, Stein TD, Alvarez VE, Daneshvar DH, et al. The spectrum of disease in chronic traumatic encephalopathy. Brain J Neurol. 2013;136:43–64.

27. Crary JF, Trojanowski JQ, Schneider JA, Abisambra JF, Abner EL, Alafuzoff I, et al. Primary age-related tauopathy (PART): a common pathology associated with human aging. Acta Neuropathol (Berl). 2014;128:755–66.

28. Pezzotti P, Scalmana S, Mastromattei A, Di Lallo D. The accuracy of the MMSE in detecting cognitive impairment when administered by general practitioners: A prospective observational study. BMC Fam Pract. 2008;9:29.

29. Fawcett T. ROC Graphs: Notes and Practical Considerations for Researchers. 2004.

30. McKenzie AT, Katsyv I, Song W-M, Wang M, Zhang B. DGCA: A comprehensive R package for Differential Gene Correlation Analysis. BMC Syst Biol [Internet]. 2016 [cited 2019 May 2];10. Available from: https://www.ncbi.nlm.nih.gov/pmc/articles/PMC5111277/

31. Gutman DA, Khalilia M, Lee S, Nalisnik M, Mullen Z, Beezley J, et al. The Digital Slide Archive: A Software Platform for Management, Integration, and Analysis of Histology for Cancer Research. Cancer Res. 2017;77:e75–8.

32. Scutari M. Learning Bayesian Networks with the bnlearn R Package. J Stat Softw. 2010;35:1–22.

33. Filley CM. Cognitive Dysfunction in White Matter Disorders: New Perspectives in Treatment and Recovery. J Neuropsychiatry Clin Neurosci. 2021;appineuropsych21030080.

34. McAleese KE, Miah M, Graham S, Hadfield GM, Walker L, Johnson M, et al. Frontal white matter lesions in Alzheimer’s disease are associated with both small vessel disease and AD-associated cortical pathology. Acta Neuropathol (Berl). 2021;142:937–50.

35. McAleese KE, Walker L, Graham S, Moya ELJ, Johnson M, Erskine D, et al. Parietal white matter lesions in Alzheimer’s disease are associated with cortical neurodegenerative pathology, but not with small vessel disease. Acta Neuropathol (Berl). 2017;134:459–73.

36. Hill RA, Li AM, Grutzendler J. Lifelong cortical myelin plasticity and age-related degeneration in the live mammalian brain. Nat Neurosci. 2018;21:683–95.

37. Marner L, Nyengaard JR, Tang Y, Pakkenberg B. Marked loss of myelinated nerve fibers in the human brain with age. J Comp Neurol. 2003;462:144–52.

38. Peters A. The effects of normal aging on myelin and nerve fibers: a review. J Neurocytol. 2002;31:581–93.

39. Alosco ML, Stein TD, Tripodis Y, Chua AS, Kowall NW, Huber BR, et al. Association of White Matter Rarefaction, Arteriolosclerosis, and Tau With Dementia in Chronic Traumatic Encephalopathy. JAMA Neurol. 2019;76:1298–308.

40. Pfefferbaum A, Adalsteinsson E, Sullivan EV. Dysmorphology and microstructural degradation of the corpus callosum: Interaction of age and alcoholism. Neurobiol Aging. 2006;27:994–1009.

41. Rosenberg GA, Wallin A, Wardlaw JM, Markus HS, Montaner J, Wolfson L, et al. Consensus statement for diagnosis of subcortical small vessel disease. J Cereb Blood Flow Metab Off J Int Soc Cereb Blood Flow Metab. 2016;36:6–25.

42. Hachinski VC, Potter P, Merskey H. Leuko-araiosis. Arch Neurol. 1987;44:21–3.

43. Marek M, Horyniecki M, Frączek M, Kluczewska E. Leukoaraiosis - new concepts and modern imaging. Pol J Radiol. 2018;83:e76–81.

44. Wardlaw JM, Smith EE, Biessels GJ, Cordonnier C, Fazekas F, Frayne R, et al. Neuroimaging standards for research into small vessel disease and its contribution to ageing and neurodegeneration. Lancet Neurol. 2013;12:822–38.

45. Pantoni L, Garcia JH. Pathogenesis of leukoaraiosis: a review. Stroke. 1997;28:652–9.

46. Haneuse S, Schildcrout J, Crane P, Sonnen J, Breitner J, Larson E. Adjustment for selection bias in observational studies with application to the analysis of autopsy data. Neuroepidemiology. 2009;32:229–39.

47. Nelson PT, Dickson DW, Trojanowski JQ, Jack CR, Boyle PA, Arfanakis K, et al. Limbic-predominant age-related TDP-43 encephalopathy (LATE): consensus working group report. Brain. 2019;142:1503–27.

48. Sekiyama K, Takamatsu Y, Koike W, Waragai M, Takenouchi T, Sugama S, et al. Insight into the Dissociation of Behavior from Histology in Synucleinopathies and in Related Neurodegenerative Diseases. J Alzheimers Dis JAD. 2016;52:831–41.

